# ERGA-BGE genome of *Valencia hispanica* (Valenciennes, 1826): a critically endangered Iberian toothcarp

**DOI:** 10.1101/2024.07.31.604920

**Authors:** Marc Ventura, Nati Franch, Rosa Fernández, Javier Palma-Guerrero, Astrid Böhne, Rita Monteiro, Laura Aguilera, Marta Gut, Tyler S. Alioto, Francisco Câmara Ferreira, Fernando Cruz, Jèssica Gómez-Garrido, Leanne Haggerty, Fergal Martin, Thomas Brown

**Affiliations:** Centre for Advanced Studies of Blanes, Spanish National Research Council (CEAB-CSIC), Accés Cala Sant Francesc 14, 17300 Blanes, Catalonia, Spain; Parc Natural del Delta de l’Ebre. Av. Catalunya, 46, 43580, Deltebre, Tarragona, Catalonia, Spain; Metazoa Phylogenomics Lab, Biodiversity Program, Institute of Evolutionary Biology (IBE, CSIC-UPF), Passeig maritim de la Barceloneta 37-49, 08003, Barcelona, Spain; FIBL Research Institute of Organic Agriculture, Ackerstrasse 113, 5070 Frick, Switzerland; Leibniz Institute for the Analysis of Biodiversity Change, Museum Koenig Bonn, Adenauerallee 127, 53113 Bonn, Germany; Centro Nacional de Análisis Genómico (CNAG), Baldiri Reixac 4, 08028 Barcelona, Spain; Universitat de Barcelona (UB), Barcelona, Spain; European Molecular Biology Laboratory, European Bioinformatics Institute, Wellcome Genome Campus, Hinxton, Cambridge, UK; Leibniz Institute for Zoo and Wildlife Research, Alfred-Kowalke-Straße 17, 10315 Berlin, Germany; Berlin Center for Genomics in Biodiversity Research (BeGenDiv), Koenigin-Luise-Str 6-8, 14195 Berlin, Germany

**Keywords:** *Valencia hispanica*, genome assembly, European Reference Genome Atlas, Biodiversity Genomic Europe, Earth Biogenome Project, Valenciidae, Valencia toothcarp, Samaruc, Samarugo

## Abstract

The reference genome of *Valencia hispanica*, a critically endangered actinopterygian species endemic to the Iberian Peninsula, is key to unravelling its genetic architecture and adaptation to freshwater ecosystems. This genomic resource will enable targeted conservation efforts and shed light on the species’ essential role in ecological dynamics, including its contributions to algal biomass regulation and role in the aquatic food web while also highlighting the challenges it faces from habitat degradation and invasive species. Furthermore, it offers opportunities to gain valuable insights into the evolutionary paths within the Valenciidae family, significantly advancing our comprehension of genetic diversity and adaptability in aquatic ecosystems. The entirety of the genome sequence was assembled into 24 contiguous chromosomal pseudomolecules. This chromosome-level assembly encompasses 1.29Gb, composed of 99 contigs and 28 scaffolds, with contig and scaffold N50 values of 38.3Mb and 56.9Mb, respectively.

## Introduction

*Valencia hispanica* (Valenciennes, 1826), commonly known as ‘samaruc’ (catalan), is an actinopterygian fish in the family Valenciidae, inhabiting euryhaline still waters locally called “ullals’’ and streams of the coast of the Mediterranean Sea. Endemic to the south of Catalonia and Valencia in the Iberian Peninsula, this species has suffered a significant reduction of its distributional range, with only ten populations remaining in the wild (Oliva-Paterna et al., 2009). It is a killifish species with slow growth compared to other Cyprinodontiformes and a restricted trophic spectrum (Rincón et al., 2002). *V. hispanica* is currently classified as “Critically Endangered” on the IUCN Red List, reflecting its high risk of extinction in the wild due to significant habitat loss and the threat by introduced exotic fish species such as Eastern mosquitofish (Caiola & de Sostoa, 2005). It is also listed under the Berne Convention on the Conservation of European Wildlife and Natural Habitats (1992/43/EEC) and as a priority species in the European Habitats Directive (under appendix II, 92/43/EEC), and in the Spanish Royal Decree 139/2011. It has a Recovery Plan in Catalonia (Catalonian Government Decree 259/2004) and Valencia (Valencian Government Decree 265/2004). Species recovery is aided by conservation initiatives by releasing captive individuals into the wild (Enric Aparicio, 2016; Oliva-Paterna et al., 2009).

Genetic studies have identified strong population subdivisions of *V. hispanica* indicating reduced gene flow among existing populations (Perdices et al., 1996). *V. hispanica* is a key endemic species in its aquatic ecosystem. Due to its relatively narrow trophic spectrum (gammarids, midges, and isopods, Rincón et al., 2002), it is vital for maintaining the ecological balance, as it contributes to controlling algal blooms and provides a key food for larger predatory fish and bird species, thus supporting the overall biodiversity and health of the ecosystem (Caiola et al., 2001).

The generation of a high-quality reference genome for *V. hispanica* will enable us to gain a deeper understanding of its genetic makeup and adaptive traits regarding global change. This genomic resource will not only aid in conservation efforts by informing breeding and habitat restoration programs but will also provide valuable insights into evolutionary processes and genetic diversity within the Valenciidae family, offering broader implications for aquatic genomic research.

The generation of this reference resource was coordinated by the European Reference Genome Atlas (ERGA) initiative’s Biodiversity Genomics Europe (BGE) project, supporting ERGA’s aims of promoting transnational cooperation to promote advances in the application of genomics technologies to protect and restore biodiversity (Mazzoni et al., 2023).

## Materials & Methods

ERGA’s sequencing strategy includes Oxford Nanopore Technology (ONT) and/or Pacific Biosciences (PacBio) for long-read sequencing, along with Hi-C sequencing for chromosomal architecture, Illumina Paired-End (PE) for polishing (i.e. recommended for ONT-only assemblies), and RNA sequencing for transcriptomic profiling, to facilitate genome assembly and annotation.

### Sample and Sampling Information

On March 22, 2023, one female specimen of *Valencia hispanica* was sampled by Marc Ventura. The identification of the species was confirmed by Nati Franch from the Ichthyological Institute in Ebro Delta, Catalonia, Spain, using the guide “Peixos continentals de Catalunya” (Aparicio, 2016). The specimen originated from a culture initially sourced from Llacunes de Santes Creus, Catalonia, Spain at the Ichthyological Center of the Ebro Delta (Ebro Delta Naltura Park). Sampling was conducted under the permit FUE-2023-03130382, issued by the Catalan Government. A fyke net was used for the collection. Specimens were euthanized by placing them on dry ice and subsequently preserved on dry ice. Until DNA extraction, the samples were stored in liquid nitrogen.

### Vouchering information

Physical reference materials for the same sequenced specimen have been deposited in Museo Nacional de Ciencias Naturales (MNCN, CSIC) https://mncn.csic.es/en under the accession number MNCN_ICTIO 296.954. Frozen reference tissue material of head, gonads, liver, and muscle is available from the same individual at the Biobank Museo Nacional de Ciencias Naturales (MNCN, CSIC) https://mncn.csic.es/en under the voucher ID MNCN-ADN-110987.

## Data Availability

*V. hispanica* and the related genomic study were assigned to Tree of Life ID (ToLID) ‘fValHis1’ and all sample, sequence, and assembly information are available under the umbrella BioProject PRJEB67580. The sample information is available at the following BioSample accessions: SAMEA113595463, SAMEA113595466, and SAMEA113595473. The genome assembly is accessible from GenBank under accession number GCA_963556495.2 and the annotated genome is available through the Ensembl Rapid Release page (https://projects.ensembl.org/erga-bge/). Sequencing data produced as part of this project are available from SRA at the following accessions: ERX11496322, ERX11496323, and ERX11496325. Documentation related to the genome assembly and curation can be found in the ERGA Assembly Report (EAR) document available at https://github.com/ERGA-consortium/EARs/blob/main/Assembly_Reports/Valencia_hispanica/fValHis1/fValHis1_EAR.pdf. Further details and data about the project are hosted on the ERGA portal at https://portal.erga-biodiversity.eu/organism/SAMEA113595463.

### Genetic Information

The estimated genome size, based on ancestral taxa, is 1.53Gb. This is a diploid genome with a haploid number of 24 chromosomes (2n=48) and unknown sex chromosomes. All information for this species was retrieved from Genomes on a Tree (Challis R et al., 2023).

### DNA/RNA processing

DNA was extracted from muscle tissue using the Blood & Cell Culture DNA Midi Kit (Qiagen) following the manufacturer’s instructions. DNA quantification was performed using a Qubit dsDNA BR Assay Kit (Thermo Fisher Scientific), and DNA integrity was assessed using a Genomic DNA 165 Kb Kit (Agilent) on the Femto Pulse system (Agilent). The DNA was stored at +4ºC until used.

RNA was extracted using an RNeasy Mini Kit (Qiagen) according to the manufacturer’s instructions. RNA was extracted from four different specimen parts: ovary, head, muscle, and liver. RNA quantification was performed using the Qubit RNA BR kit and RNA integrity was assessed using a Bioanalyzer 2100 system (Agilent) RNA 6000 Pico Kit (Agilent). RNA was pooled equimolarly for the library preparation and stored at −80ºC until used.

### Library Preparation and Sequencing

For the long-read whole sequencing, the library was prepared using the SQK-LSK114 Kit (Oxford Nanopore Technologies, ONT), and the library was sequenced on PromethION 24 A Series instrument (ONT). A short-read whole genome sequencing library was prepared using the KAPA Hyper Prep Kit (Roche). Hi-C library was prepared from liver tissue using the Dovetail Omni-C kit (Cantata Bio). The RNA library from the pooled sample was prepared using the KAPA mRNA Hyper prep kit for Illumina sequencing (Roche). The short-read sequencing libraries were processed on a NovaSeq 6000 instrument (Illumina). In total 74X Oxford Nanopore, 83X Illumina WGS shotgun, and 70X HiC data were sequenced to generate the assembly.

### Genome Assembly Methods

The genome was assembled using the CNAG CLAWS pipeline (Jessica Gomez-Garrido, 2024). Briefly, reads were preprocessed for quality and length using Trim Galore v0.6.7 and Filtlong v0.2.1, and initial contigs were assembled using NextDenovo v2.5.0, followed by polishing of the assembled contigs using HyPo v1.0.3, removal of retained haplotigs using purge-dups v1.2.6 and scaffolding with YaHS v1.2a. Finally, assembled scaffolds were curated via manual inspection using Pretext v0.2.5 with the Rapid Curation Toolkit (https://gitlab.com/wtsi-grit/rapid-curation) to remove any false joins and incorporate any sequences not automatically scaffolded into their respective locations in the chromosomal pseudomolecules (or super-scaffolds). Finally, the mitochondrial genome was assembled as a single circular contig of 16,558 bp using the FOAM pipeline (https://github.com/cnag-aat/FOAM) and included in the released assembly (GCA_963556495.2).

### Genome Annotation Methods

A gene set was generated using the Ensembl Gene Annotation system (Aken et al., 2016), primarily by aligning publicly available short-read RNA-seq data from BioSample: SAMEA113595463 to the genome. Gaps in the annotation were filled via protein-to-genome alignments of a select set of vertebrate proteins from UniProt (The UniProt Consortium, 2019), which had experimental evidence at the protein or transcript level. At each locus, data were aggregated and consolidated, prioritising models derived from RNA-seq data, resulting in a final set of gene models and associated non-redundant transcript sets. To distinguish true isoforms from fragments, the likelihood of each open reading frame (ORF) was evaluated against known vertebrate proteins. Low-quality transcript models, such as those showing evidence of fragmented ORFs, were removed (thresholds needed). In cases where RNA-seq data were fragmented or absent, homology data were prioritised, favouring longer transcripts with strong intron support from short-read data. The resulting gene models were classified into three categories: protein-coding, pseudogene, and long non-coding. Models with hits to known proteins and few structural abnormalities were classified as protein-coding. Models with hits to known proteins but displaying abnormalities, such as the absence of a start codon, non-canonical splicing, unusually small intron structures (<75 bp), or excessive repeat coverage, were reclassified as pseudogenes. Single-exon models with a corresponding multi-exon copy elsewhere in the genome were classified as processed (retrotransposed) pseudogenes. Models that did not fit any of the previously described categories did not overlap protein-coding genes, and were constructed from transcriptomic data were considered potential lncRNAs. Potential lncRNAs were further filtered to remove single-exon loci due to their unreliability. Putative miRNAs were predicted by performing a BLAST search of miRBase (Kozomara et al., 2019) against the genome, followed by RNAfold analysis (Gruber et al., 2008). Other small non-coding loci were identified by scanning the genome with Rfam (Kalvari et al., 2018) and passing the results through Infernal (Nawrocki and Eddy, 2013).

## Results

### Genome Assembly

The genome assembly has a total length of 1,291,697,043 bp in 29 scaffolds including the mitogenome (Figures 1-3), with a GC content of 40.09%. The assembly has a contig N50 of 38,296,799bp and L50 of 14 and a scaffold N50 of 56,937,854bp and L50 of 11. The assembly has a total of 71 gaps, totaling 14kb in cumulative size. The single-copy gene content analysis using the Actinopterygii database with BUSCO (Manni et al., 2021) resulted in 98.6% completeness (97.9% single and 0.7% duplicated). 98.05% of reads k-mers were present in the assembly and the assembly has a base accuracy Quality Value (QV) of 50.60 as calculated by Merqury (Rhie et al., 2020).

**Figure 1.**
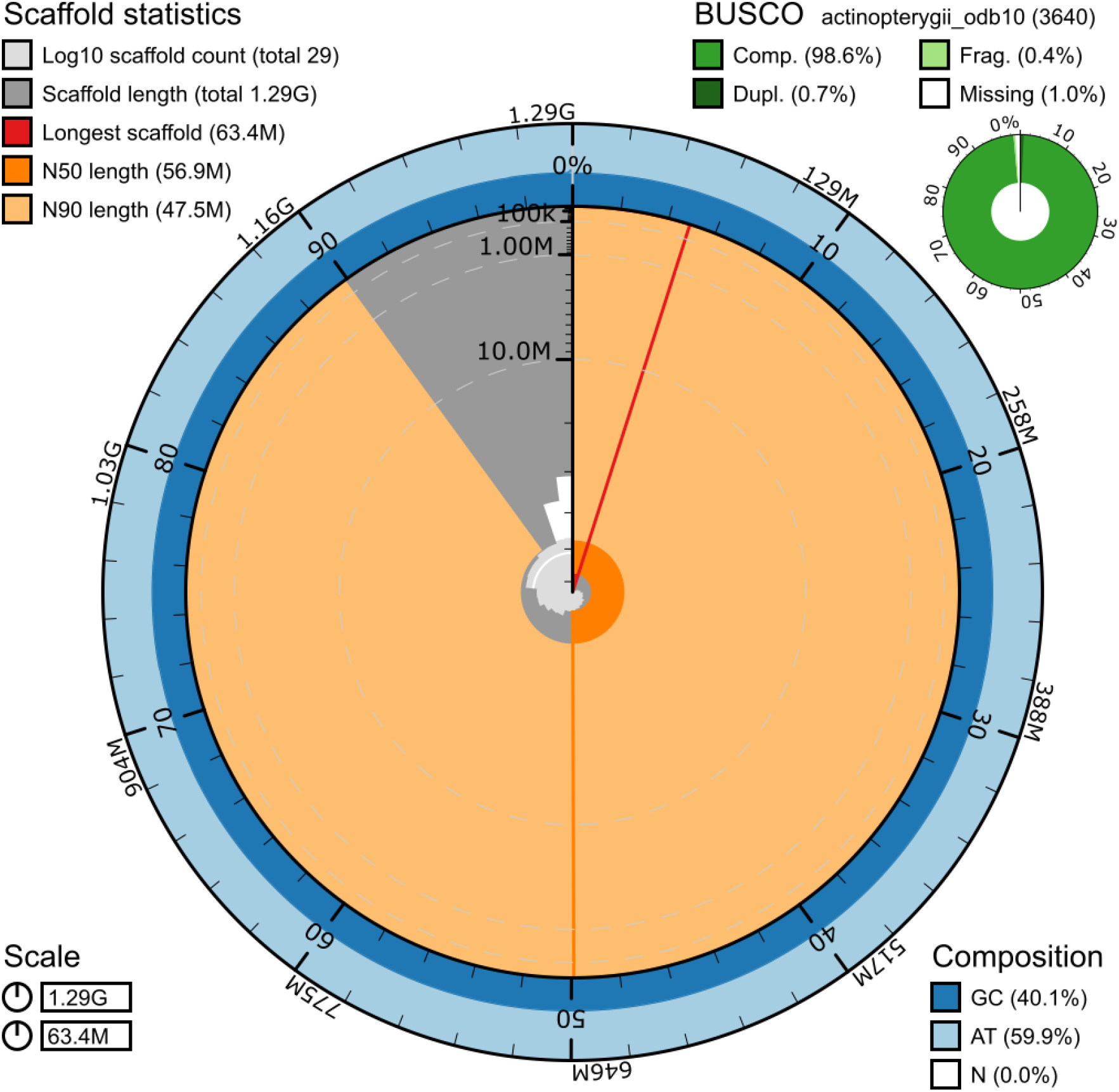
Snail plot summary of assembly statistics. The main plot is divided into 1,000 size-ordered bins around the circumference, with each bin representing 0.1% of the 1,291,697,043 bp assembly including the mitochondrial genome. The distribution of sequence lengths is shown in dark grey, with the plot radius scaled to the longest sequence present in the assembly (63,361,188 bp, shown in red). Orange and pale-orange arcs show the scaffold N50 and N90 sequence lengths (56,937,854 and 47,505,306 bp), respectively. The pale grey spiral shows the cumulative sequence count on a log-scale, with white scale lines showing successive orders of magnitude. The blue and pale-blue area around the outside of the plot shows the distribution of GC, AT, and N percentages in the same bins as the inner plot. A summary of complete, fragmented, duplicated, and missing BUSCO genes found in the assembled genome from the Actinopterygii database (odb10) is shown in the top right.

**Figure 2.**
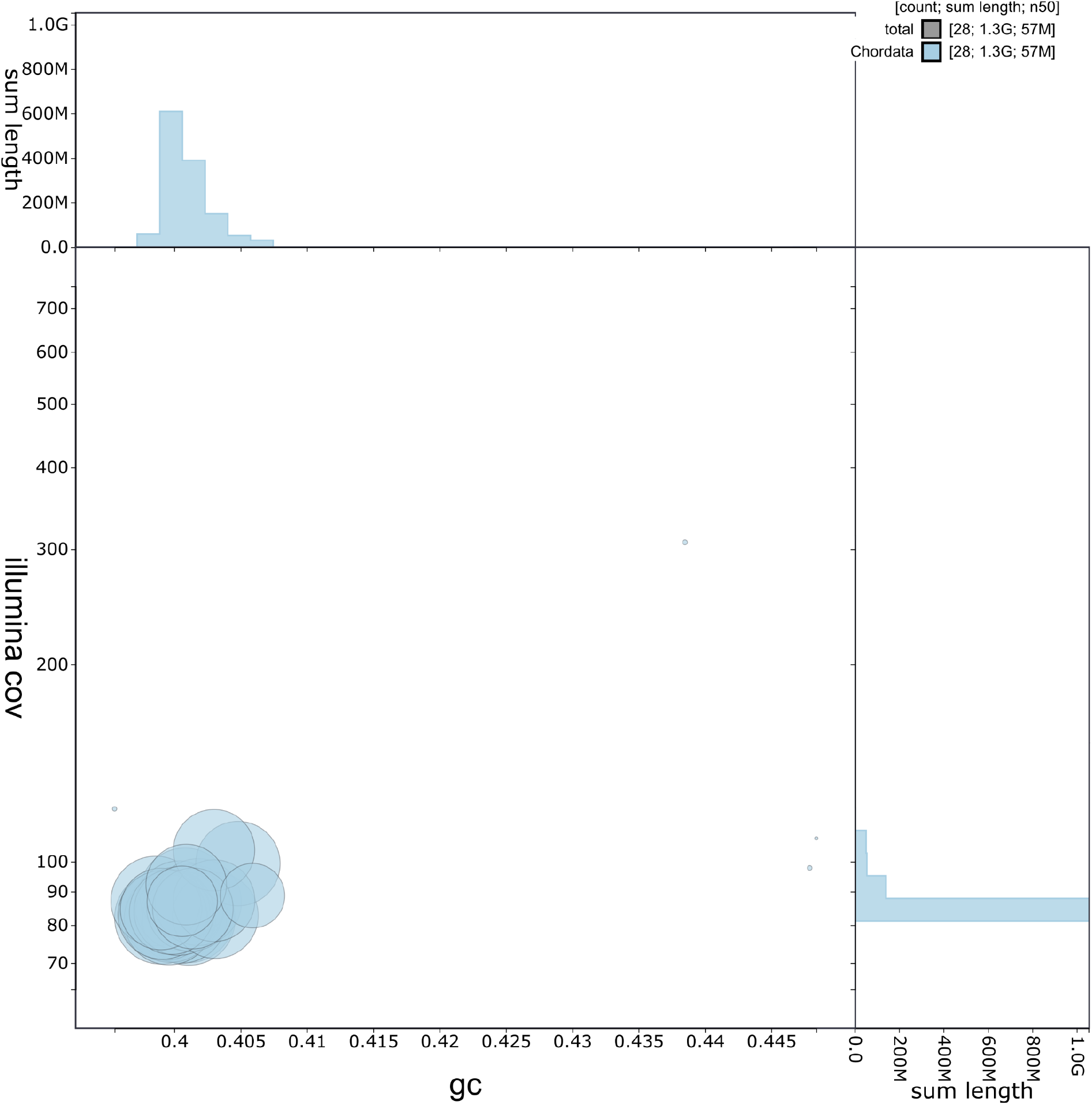
Blob plot contamination screening showing coverage of Illumina WGS reads against GC proportion per scaffold in the assembly. Sequences are coloured by phylum. Blob size is proportional to the sequence length. Histograms show the distribution of sequence length sum along each axis.

**Figure 3.**
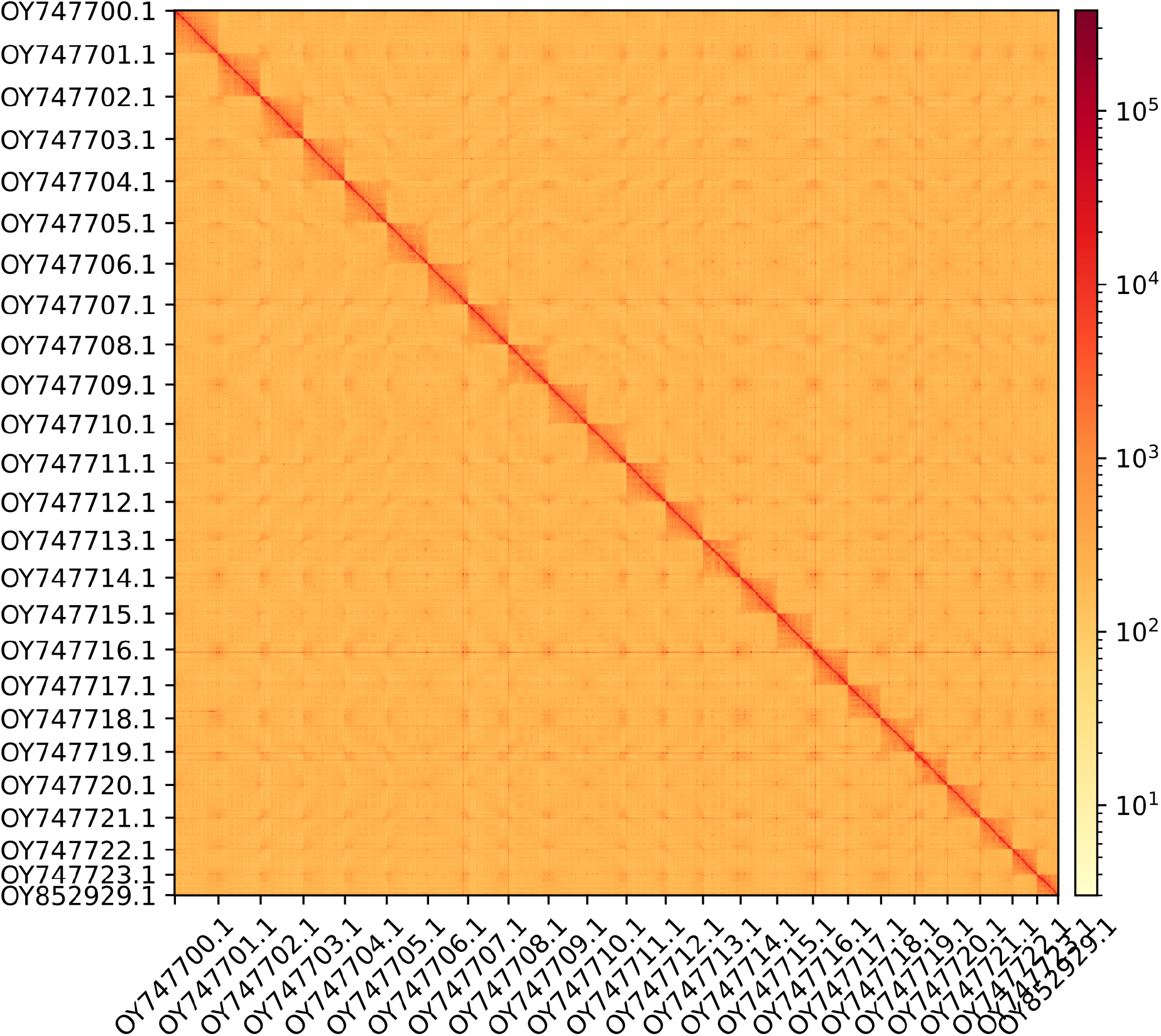
Hi-C contact map showing spatial interactions between regions of the genome. The diagonal corresponds to intra-chromosomal contacts, depicting chromosome boundaries. The frequency of contacts is shown on a logarithmic heatmap scale. Hi-C matrix bins were merged into a 100 kb bin size for plotting.

### Genome Annotation

The genome annotation consists of 20,447 protein-coding genes with an associated 32,953 transcripts, in addition to 1,600 non-coding genes (Table 1). Using the longest isoform per transcript, the single-copy gene content analysis using the Actinopterygii database with BUSCO resulted in 94.2% completeness. Using the OMAmer Cyprinodontoidei database for OMArk (Nevers et al., 2024) resulted in 91.54% completeness and 98.39% consistency (Table 2)

**Table 1.** Statistics from assembled gene models.

|  | No. genes | No. transcripts | Mean gene length (bp) | No. single-exon genes | Mean exons per transcript |
| --- | --- | --- | --- | --- | --- |
| mRNA | 20,447 | 32,953 | 26,493 | 517 | 11.5 |
| pseudogene | 71 | 71 | 42,642 | 2 | 21.2 |
| snoRNA | 184 | 184 | 123 | 184 | 1 |
| lncRNA | 271 | 193 | 11,288 | 11 | 3.7 |
| miRNA | 39 | 39 | 69 | 39 | 1 |
| snRNA | 310 | 310 | 147 | 310 | 1 |
| rRNA | 668 | 668 | 611 | 668 | 1 |
| scRNA | 7 | 7 | 226 | 7 | 1 |
| Other ncRNA | 50 | 50 | 298 | 50 | 1 |

**Table 2.** Annotation completeness and consistency scores calculated by BUSCO run in protein mode (Actinopterygii_odb10) and OMArk (Cyprinodontoidei)

|  | Complete | Singular | Duplicated | Fragmented | Missing |
| --- | --- | --- | --- | --- | --- |
| BUSCO | 3,428 (94.2%) | 3382 (92.9%) | 46 (1.3%) | 40 (1.1%) | 172 (4.7%) |
| OMArk | 16,981 (91.54%) | 16,510 (89.53%) | 371 (2.01%) | - | 1,560 (8.46%) |
|  | Consistent | Inconsistent | Contaminants | Unknown |  |
| OMArk | 32,441 (98.39%) | 387 (1.17%) | 0 (0.00%) | 143 (0.43%) |  |

## Acknowledgements

We would like to thank the Ebro Delta Ichthyological Center from the Ebro Delta Natural Park (Catalan Government) and their personnel for maintaining a stock population of *Valencia hispanica*, performing monitoring and conservation works, and providing the individuals for sequencing the genome (specific permit FUE-2023-03130382). The restoration projects in Catalonia (e.g. SOS Samaruc) are funded by Fundació Andrena with contributions from Agència Catalana de l’Aigua (Catalan Government) and Fundación Biodiversidad (Spanish Government) and are executed in collaboration with Associació Paissatges Vius, Ebro Delta Natural Park and Sorelló. We would like to acknowledge the assembly reviewer, Diego De Panis, from the Leibniz Institute for Zoo and Wildlife Research.

## Author Contributions

JPG and RF coordinated the project, MV collected the species, NF identified the species, MV sampled and preserved biological material and provided metadata, AB and RM provided sampling and metadata support and management, LA and MG extracted DNA, prepared libraries, and performed sequencing, FCF JGG and FC performed genome assembly and curation under the supervision of TA, LH, and FM performed genome annotation, TB generated the analysis and report. All authors contributed to the writing, review, and editing of this genome note and read and approved the final version.

